# 3D chromatin structure estimation through a constraint-enhanced score function

**DOI:** 10.1101/075184

**Authors:** Claudia Caudai, Emanuele Salerno, Monica Zoppè, Anna Tonazzini

## Abstract

Based on experimental techniques of the type *Chromosome Conformation Capture* (3C), several methods have been proposed in the literature to estimate the structure of the nuclear dna in homogeneous populations of cells. Many of these methods transform contact frequencies into Euclidean distances between pairs of chromatin fragments, and then reconstruct the structure by solving a distance-to-geometry problem. To avoid the drawbacks of this strategy, we propose to abandon the frequency-distance translation and adopt a recursive multiscale procedure, where the chromatin fibre is modelled by a new kind of modified bead chain, the data are suitably partitioned at each scale, and the resulting partial structures are estimated independently of each other and then connected again to rebuild the whole chain.

We propose a new score function to generate the solution space: it includes a data-fit part that does not require target distances, and a penalty part, which enforces *soft* geometric constraints on the solution, coherent with known physical and biological constraints. The relative weights of the two parts are balanced automatically at each scale and each subchain treated. Since it is reasonable to expect that many different structures fit any 3c-type data set, we sample the solution space by simulated annealing, with no search for an absolute optimum. A set of different solutions with similar scores is thus generated. The procedure can be managed through a minimum set of parameters, independent of both the scale and the particular genomic segment being treated. The user is thus allowed to control the solutions easily and effectively. The partition of the fibre, along with several intrinsically parallel parts, make this method computationally efficient.

We report some results obtained with the new method and code, tested against real data, that support the reliability of our method and the biological plausibility of our solutions.

## Background

*Chromosome Conformation Capture* (3c, [1]) and a number of related techniques (4c, [2], 5c, [3], Hi-C, [4, 5]) provide high-throughput, high-resolution matrices of contact frequencies between pairs of dna fragments in uniform populations of cells. A number of strategies have been proposed in recent literature to use these data to estimate the spatial structures possibly assumed by the chromatin fibre inside the cell nuclei. The proposed approaches vary in chromatin fibre modelling and in the sampling strategy of the solution space. Irrespective of the type of approach, some of the most recent proposals are in [6–9]. A more comprehensive and reasoned panorama can be drawn, for example, from [10–14]. Many of these methods use an explicit relationship between the contact frequencies and the Euclidean distances between all the possible pairs of dna fragments. The sensible motivation of this choice is that pairs of fragments that are frequently in contact are likely to be spatially close, whereas pairs of fragments with a few contacts are supposed to be farther apart.

After a critical analysis of the problem, we pose three main desirable requirements for an estimation algorithm. First of all, as the number of possible pairs in any real-world case is enormous, the algorithm must be computationally efficient, allowing partitioning strategies and parallel processing to be applied. Second, since the data are produced in experiments on millions of cells, very different structures are likely to contribute in the contact frequency matrix, so the algorithm must be capable of estimating different configurations compatible with the data. For a discussion on this point, see [11] or [13]. Last, but not least, translating contact frequencies into Euclidean distances does not seem appropriate [14–16], since two fragments that are seldom in contact can well be spatially close. Any target distance derived directly from the frequency data should thus be avoided. Moreover, the algorithm should offer the possibility of incorporating known geometrical constraints into the solutions.

The estimation method we propose is characterised by a multiresolution, modified bead-chain model for the chromatin fibre. We motivate this choice from the observation that the contact matrices typically highlight genomic regions characterised by many internal contacts and poor interactions with the rest of the genome. This feature is found practically at all the scales [11, 17]: at 1-10 Mbp scale, the *genomic compartments* show this property; at smaller scales (100 kbp and below), structures that behave similarly are referred to as *topological association domains* (or TADS).

Taking this recursive behaviour into account, our strategy is to identify high-interaction diagonal blocks in the original data matrix. Since the related chromatin segments interact very weakly with the rest of the genome, their structures can be estimated irrespective of the rest of the matrix. Then, each estimated segment becomes part of a lower-resolution chain as a modified bead only characterised by its endpoints and its centroid. After an appropriate binning of the data matrix, the lower-resolution chain is further partitioned and reconstructed, and its resolution is lowered, recursively, until the binned matrix cannot be partitioned anymore. In our approach, the fit to the data is not measured through target distances, but through the closeness of the most frequently interacting pairs of beads. All the other beads are free to follow the likely configurations of the chromatin chain. A score function based on this measure is built to generate a solution space that is then sampled to estimate the likely structures at the various scales, subject to adequate constraints so as to avoid the physically unfeasible solutions. A number of configurations is thus generated, compatible with both the data and their expected geometrical features. Once the solutions at all scales are available, a backward recursive procedure aligns the subchains reconstructed with the corresponding lower-resolution beads, repeating until all the subchains at full resolution are properly located and aligned.

An algorithm implementing this strategy is presented in [14], where the geometric constraints consist in minimum allowed mutual distances between beads and maximum allowed planar angles between consecutive triples of beads. These constraints are imposed rigidly: any solution that violates them is simply discarded from the sampling. The solution space is sampled by simulated annealing, and the proposed structures are evolved through quaternion operators [18].

In this paper, we propose a method that follows the same general strategy, but uses a new score function including implicit constraints and a unified bead model at all the scales. Rather than simply rejecting the unfeasible solutions, the implicit constraints penalise gradually the unlikely configurations, and the data-fit term favours the configurations where the closest pairs of fragments are those with the highest contact frequencies. The free parameters to be set are very few, and this allows us to control the solutions easily and effectively. The code implementing this method, now at its version 3.1, is written in Python and is called ChromStruct, available online^1^ in its command-line and GUI versions, with an example data set, a separate visualisation code and some explanatory notes.

In the Methods section, we give the details of the solution model, the score function, and the recursive procedure. In the Results and discussion section we analyse the performance of ChromStruct 3.1, the geometrical features of a set of results obtained using real data, and show how some of these features are correlated with known biological properties. Final remarks on the current results and possible future directions are given in the Conclusion section.

## Methods

### Chromatin model

As explained above, approximate diagonal blocks are extracted from the data matrix by a method suggested in [11]. Each block maps the contacts within a subchain, whose structure is then estimated. At the end of this process, each sub-chain becomes a single bead in a lower-resolution chain, whose internal contacts sum up to the total contacts in the corresponding block, and become a single entry of the data matrix at the successive resolution. The bead is characterised by its centroid, its endpoints, and its approximate size.

At the maximum available resolution, that is, at the first recursion level, the bead structures are not known; in this framework, this means that the first-level beads can only be modelled as spheres, whose radii can be approximated from the number of their internal contacts. Indeed, intuitively, many internal contacts mean that a fragment is packed tightly, whereas a few contacts mean that it is relatively stretched out. The treatment, however, is unified: each first-level spherical bead is modelled like the triples at the successive levels (endpoint 1; centroid; endpoint 2), but the three points are collinear, and the distances between the centroid and the two endpoints are both equal to the estimated radius. The size of a lower-resolution bead is estimated as a fraction of the maximum span of the related subchain, obtained as the strength of its first principal component. The beads are linked respecting their biological order: the second endpoint of each bead and the first endpoint of the successive bead coincide. Figure 1 illustrates how four consecutive subchains are schematised as modified beads and then connected to form a chain at a lower resolution. The lengths of the segments joining the endpoints with the centroid, and the related angle, are not changed during the evolution of the model. Conversely (see also [14]), the planar and dihedral angles defining the position of each bead with respect to the adjacent ones are perturbed at each iteration, subject to constraints that establish chain flexibility and mutual distance ranges. Our score function allows us to avoid special constraints on angles, as the bead sizes alone provide a good control.

**Figure1.**
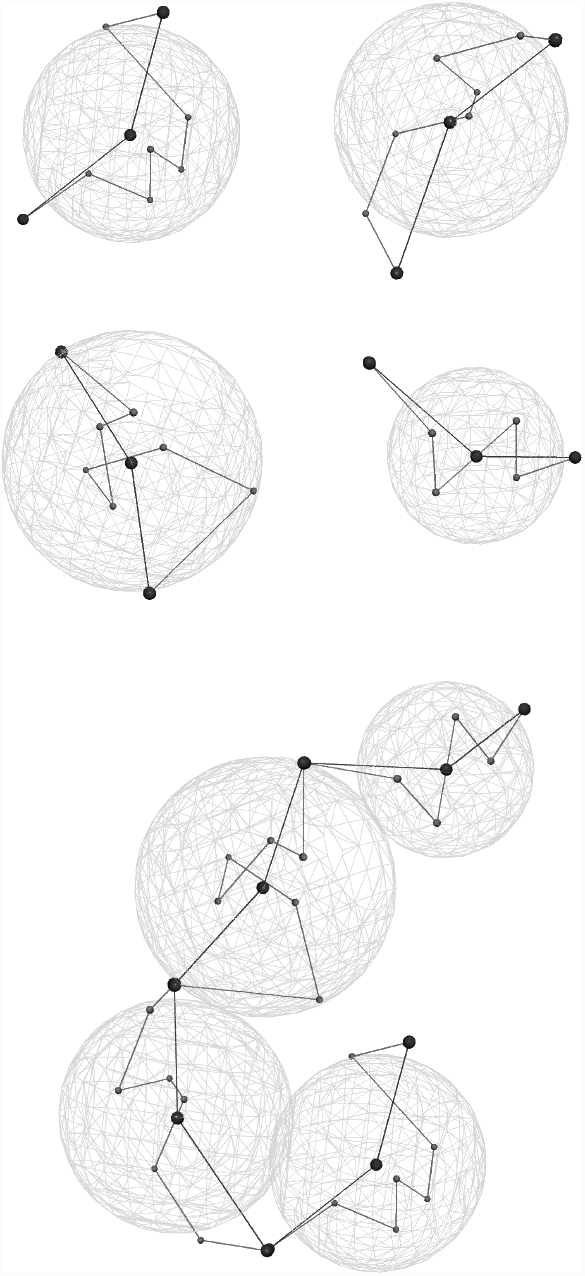
Illustrative picture of the modified bead-chain model. Top: Four chromatin fragments, represented as bead sequences (small red spheres), and as centroid-endpoints triples (bigger blue spheres). The green sphere wireframes represent the assumed sizes for the beads at the lower resolution. Bottom: Connected chain composed by the fragments above, properly located and rotated. The sequence of green spheres represents the lower-resolution chain.

### Score function

The solution space for each subchain to be estimated is generated by a function of its configuration 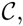, with the following form:

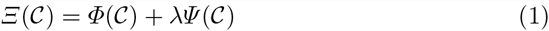

where *Φ* and *Ψ* are the data-fit and the constraint terms, respectively, and *λ* is a parameter that balances their influence. The two terms are sums of positive contributions, *ϕ*_*ij*_ and *ψ*_*ij*_, intended to penalise unlikely interactions between the beads indexed by *i* and *j*.

As far as data fit is concerned, in the light of the considerations exposed in the Background section, we make *ϕ*_*ij*_ proportional to the squared distance between the *i*-th and the *j*-th beads, but we limit the summation to a suitably chosen subset 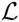 of highly interacting pairs in 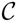:

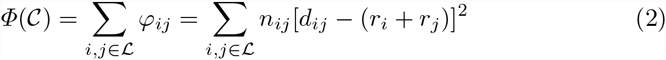

where *n*_*ij*_ is the contact frequency of the *i*-th and *j*-th beads, *d*_*ij*_ is the distance between their centroids, and *r*_*i*_ and *r*_*j*_ are their radii. To build 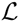 for each sub-chain, ChromStruct 3.1 selects the pairs exceeding a pre-defined percentile of the contact frequencies in the related block. Function *Φ* becomes small when the centroid-centroid distances of the pairs in 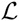 nearly equal the sums of the radii of the related beads, and grows quadratically as soon as they become smaller or larger than that values. When any two beads in 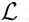 interpenetrate, one of the terms in brackets becomes negative. The maximum data-fit penalisation of this situation occurs when *d*_*ij*_ = 0, and *ϕ*_*ij*_ assumes the finite and unmodifiable value *n*_*ij*_(*r*_*i*_ + *r*_*j*_)^2^. The constraint term *Ψ* is needed to control this penalisation and extend it to all the pairs in 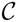. We set *Ψ* as

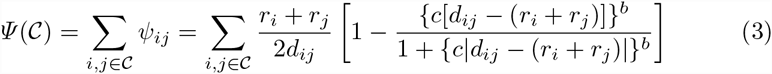

where *c* is a scale factor that makes the terms in braces dimensionless, and the exponent *b* is an odd natural. A plot of *ψ*_*ij*_ is shown in Fig. 2. Note that, for *d*_*ij*_ near zero, *ψ*_*ij*_ behaves as (*r*_*i*_ + *r*_*j*_)/*d*_*ij*_, whereas in an interval around (*r*_*i*_ +*r*_*j*_) it behaves as (*r*_*i*_ +*r*_*j*_)/(2*d*_*ij*_) and, for *d*_*ij*_ sufficiently larger than (*r*_*i*_ +*r*_*j*_), it goes rapidly to zero. Approximately, *c* tunes the width of the intermediate interval: increasing *c* means making it narrower. Parameter *b*, in turn, tunes the slope of the transitions between the different zones; large values of *b* produce abrupt transitions. Function (3) is thus intended to prevent any two beads from interpenetrating more than some fraction of their sizes. Since the bead structures are always known approximately, imposing this requirement rigidly could exclude good configurations from the feasible solutions, besides making more difficult to enforce the constraints coherently at the different scales. The position of each bead in a pair is penalised gradually as a function of its distance from the other bead. As moderate interpenetrations between adjacent beads are allowed, provided that *c* and *b* are chosen wisely, constraining the mutual angles between adjacent beads can become unnecessary.

**Figure2.**
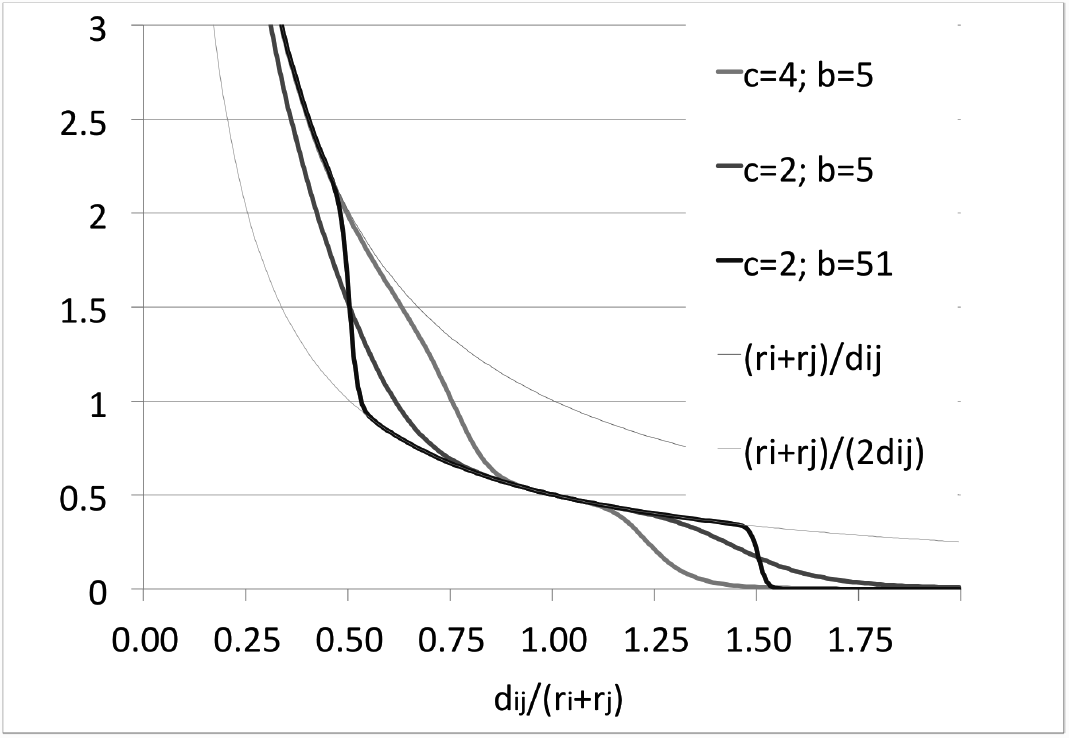
Function *ψ*_*ij*_ in (3), plotted for a few values of *c* and *b*. Increasing *c* narrows the moderate-slope interval around *d*_*i,j*_ = *r*_*i*_ + *r*_*j*_; increasing *b* increases the slopes in the two transition regions. The two thin lines represent the hyperbolas that bound the significantly nonzero values of *ψ*_*ij*_.

As said, the role of *λ* in (1) is to balance the two terms so to obtain solutions that are consistent with both the data and our prior knowledge. Its value is chosen so that, statistically, the influence of the two penalty terms is fixed in all the subchains being estimated. In our case, we have an additional difficulty related to the multiresolution setting. The extracted diagonal blocks have different sizes and frequency values, so the values assumed by the data fit term (2) for different blocks range in a very broad interval, and a unique value for *λ* working well for all the blocks cannot be established. At different scales, the situation is even worse: the orders of magnitude of *n*_*ij*_ are very diverse. To avoid the need of fixing different values *a priori*, for each block, we compute *λ* by enforcing a pre-defined ratio between the average values assumed by the two terms in (1) during a dedicated random sampling (see the source codes in the material available online).

### Estimation strategy

The solution space is sampled through an approximate simulated annealing [19], with a preliminary random sampling to estimate *λ* from the desired relative influence of the two penalty terms, and a warm-up phase to determine the start temperature for each block treated. A slow cooling schedule is then followed to reach comparable final scores in different runs.

This high level, simplified pseudocode describes the overall recursive method:

*Chromstruct* (`matrix`, `chain`, `scale`):

1. *extract diagonal blocks from* `matrix`
2. *initialise lower-resolution chain* `lo-res_chain`← null
3. For all the blocks
  a. *populate set* 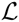;
  b. *set initial guess*: 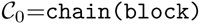
  c. *sample the penalty landscape*: 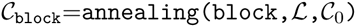
  d. *save* 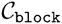
  e. *compute the equivalent low-resolution bead*: **Bead**_block_ ← *beadify*(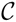)
  f. *Append* **Bead**_block_ *to* `lo-res chain`
4. *if* # *of extracted blocks* = 1 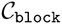 ← recursive composition of all saved 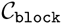 *save* 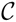 *leave* *else* *update* `scale` *bin* **matrix** *to the new resolution chromstruct*(**matrix, lo-res chain, scale**)

Since, at each scale level, each subchain is estimated independently of the others, steps 3a-e) can be performed in parallel for all the blocks. Moreover, sampling low-dimensional spaces is much less expensive than sampling the entire panorama at once. Thus, this strategy is computationally efficient. The above procedure produces *one* final structure per run. Different runs normally produce different structures. As done in [14], we could also proceed by using all the stable subchain configurations up to some scale to produce the structures at the subsequent scales. This strategy can further reduce the computing effort, and provide a means to generate many configurations from limited sets of high-level beads.

## Results and discussion

ChromStruct 3.1 was tested against real Hi-C data from human lymphoblastoid cells, chromosome 1, *q* range [150.28 Mbp, 179.44 Mbp] [4]. The original genomic resolution is 100 kbp. For these data, our code identifies two modelling levels: the first includes 292 beads, each spanning 100 kbp, and the second includes 23 beads, with genomic sizes ranging from 0.7 to 2.2 Mbp.^2^ The structures of the original and the binned matrices are visualised in Fig. 3.

**Figure3.**
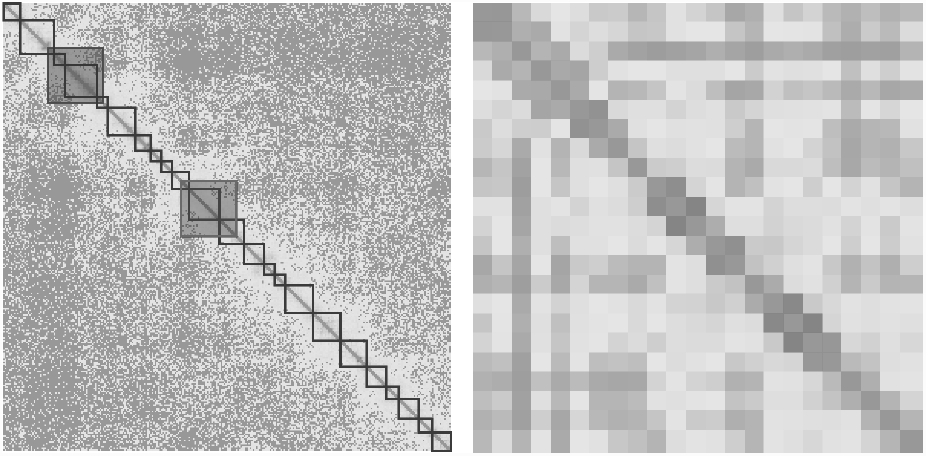
Heat maps of the contact matrix for the long arm of chromosome 1, from a HI-C experiment on human lymphoblastoid cells (GM06990, [4]). Main diagonal removed for visualisation convenience. Left: Original 292*×*292 matrix at a resolution of 100 kbp. The 23 highlighted diagonal blocks mark the topological domains used for binning. The two bigger blocks highlighted in blue (the leftmost one) and red, respectively, are related to a high-expression and a low-expression regions identified in the sequence at hand (see Biological plausibility). Right: 23 *×* 23 matrix obtained by binning the original in accordance with the extracted blocks.

We performed a first series of experiments aimed at establishing the most appropriate constraint parameters to obtain physically plausible results. All the subsequent experiments were performed with a fixed set of geometrical parameters. Their values are reported as defaults in the code provided online.

We produced 200 different solutions, from which we have first checked the reliability of the code. Next, we evaluated the relevant geometrical features of the estimated structures and looked for a possible validation from known biological properties. We report and comment these aspects in the following subsections.

### Repeatability - Fit to the data

As mentioned above, there is no unique solution optimising the score function, since the contact data come from millions of cells with different chromatin configurations. At present, thus, the only way to check the repeatability of our procedure is to ensure that the final values of the score function, for different solutions, lie in a limited range around some average value. Of course, this cannot be the same for different subchains, as their genomic sizes are different, and the contact data for different pairs are diverse. Both for each subchain and for the whole chain, however, we were able to get highly stable final scores. Through Shapiro-Francia tests [20], we also observed that all these values are compatible with Gaussian distributions.

As the data are not related to a single configuration, the data fit for any individual solution cannot be checked. However, all the single-configuration binary contact matrices can be co-added to reconstruct a synthetic contact matrix, to be compared to the input matrix. For any sufficiently numerous and diverse solution set, thus, the data fit can be appreciated. As far as diversity is concerned, we should be sure that the output structures are such that all the regions of the contact matrix are sufficiently covered [13]. One way to increase diversity is by tuning the tolerance of the annealing stop criterion. In our implementation, annealing stops when the score relative variation in 500 consecutive cycles falls below the required tolerance.

For the whole chain, our solution set was built by using tolerances of 10^*−*5^ and 10^*−*2^, obtaining normally distributed final scores with standard deviations of, respectively, 11% and 30% of the mean values. The resulting contact matrix, assuming that two beads are in contact when their distance is 1.2 times the sum of their radii,^3^ is shown in Fig. 4. From a comparison with the map in Fig. 3, it can be noted that most of the relevant structures in the original data are captured by our solutions, despite the limited number of configurations used to build the simulated matrix.

**Figure4.**
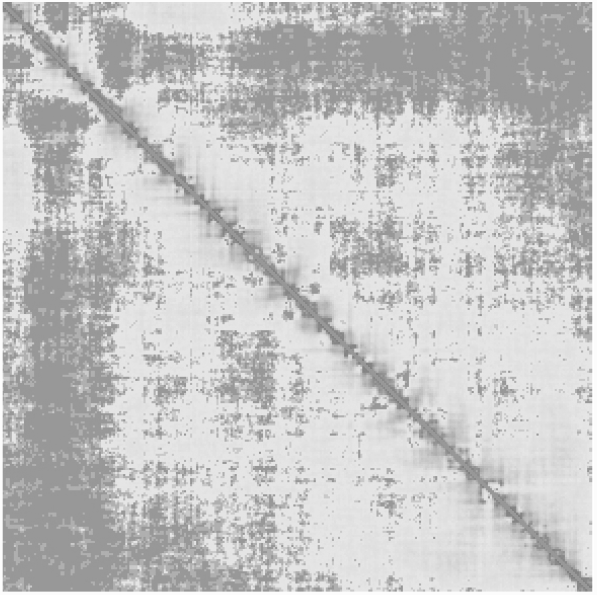
Heat map of the synthetic contact matrix built from 200 final configurations, whole chain at 100 kbp resolution, to be compared to Fig. 3, left panel.

### Geometrical features - Biological plausibility

We are interested to see whether a limited number of typical classes of configurations can be identified among our estimated structures. To this end, we clustered the results to find similarities and differences between the typical cluster members. The clustering feature we chose is the mean-squared Euclidean distance between pairs of beads as a function of their genomic distance [14, 21]. This can be indicative of the packing of each solution. The average plot of this function, evaluated from all our whole-chain configurations, is reported in Fig. 5. Considering that a fully stretched chain would produce a quadratic behaviour, this shows that most output configurations are tightly packed.

**Figure5.**
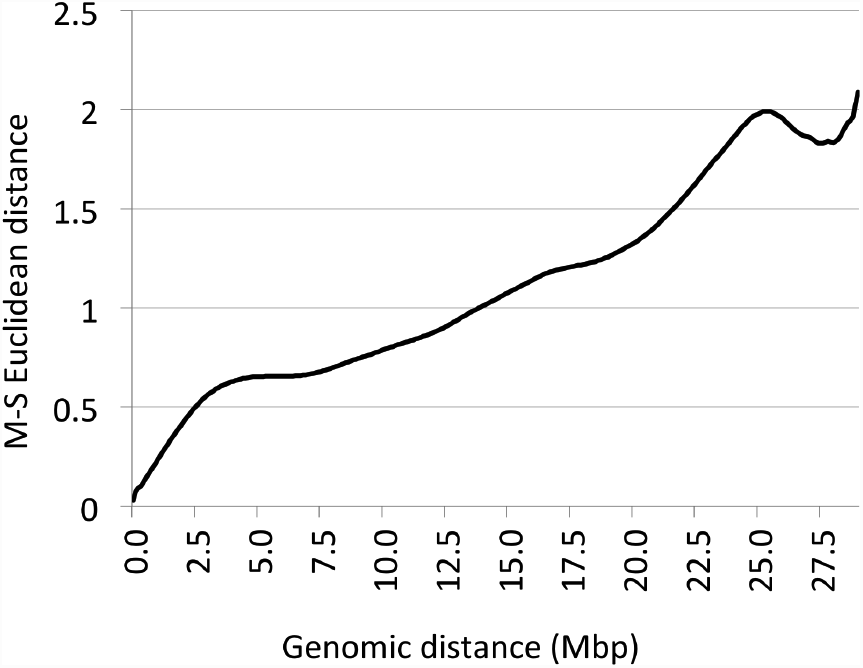
M-S Euclidean (*µ*m^2^) vs. Genomic distance plot. For each possible genomic distance in our chain, the mean-square Euclidean distances are first computed for each output configuration, and then averaged over all our 200 configurations.

To cluster the plots resulting from all the output configurations, we used the hierarchical clustering on principal components algorithm [22] (hcpc), a hybrid technique combining principal components, hierarchical, and partitional clustering, capable of detecting the number of relevant clusters from intra- and inter-cluster distance optimisation. A summary of the results of this clustering, for the whole chain, is reported in Fig. 6. Some differences between the clusters can already be detected by visual inspection.

**Figure6.**
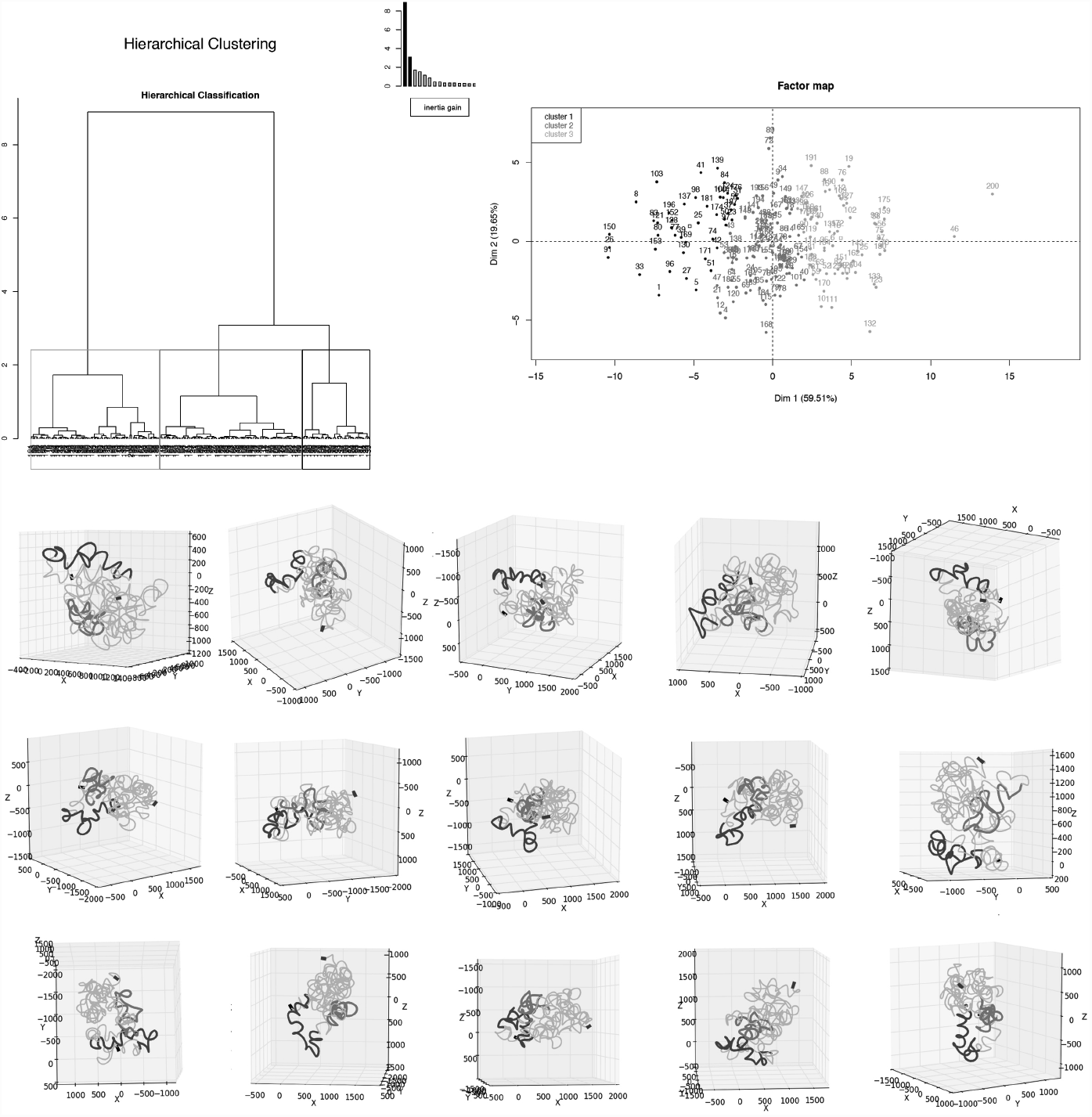
Summary of clustering results. Top: the classification tree produced by HCPC, and the three clusters detected, projected onto the first two principal components. Each of the three subsequent rows features the five configurations closest to the centroid of each cluster. The black marks locate bead 1, and the purple marks locate bead 292. Measurements in nm. The subchains highlighted in blue and red are, respectively, the highly and poorly expressed regions identified to check biological plausibility (see text, and the corresponding contact data in Fig. 3, left panel).

To check the biological plausibility of our results, we first observe that the overall size of our solutions is compatible with the biological knowledge. As this depends on the sizes of the elementary beads and the degree of packing of the chromatin fibre, this is a first index of reliability. Indeed, note that we do not scale the results *a posteriori*, but our solutions come directly with their physical dimensions.

Then, we rely on a further biological property: highly expressed or gene-rich domains are much less packed than the domains poor in genes or with low transcriptional activity [23]. Our experimental data were obtained from the human lymphoblastoid cell line GM06990, belonging to the lineage of B cells. From expression data relative to immature B cells,^4^ we identified two stretches of about 3.5 Mbp as representative of, respectively, highly expressed and poorly expressed genomic domains: dna from *q* = 151.5 Mbp to *q* = 155.1 Mbp, and dna from *q* = 162 Mbp to *q* = 165.5 Mbp. Both these regions correspond to 35 beads in our full-resolution model, and are also rich and poor in genes, respectively.^5^ Their approximate locations are highlighted by different colours in the final configurations of Fig. 6.

**Figure7.**
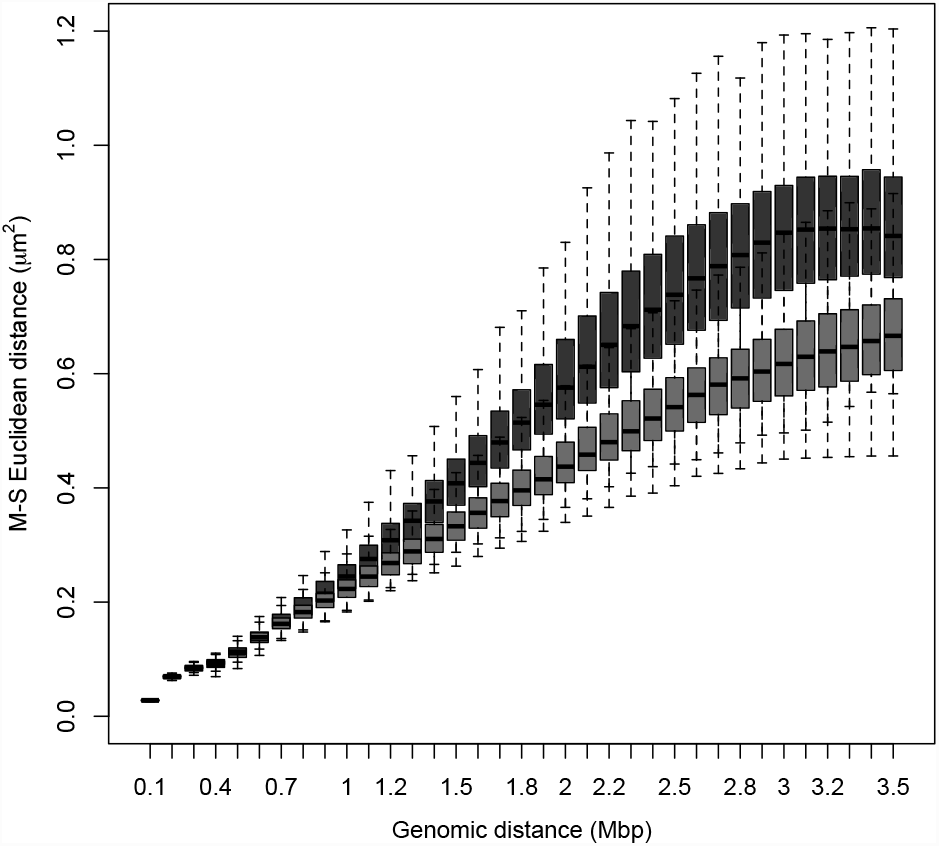
Boxplots of M-S Euclidean vs. Genomic distance, obtained from all our 200 solutions for the identified highly-expressed (blue) and poorly expressed (red) regions.

To check the packing of the corresponding segments in our estimated chains, we use again the mean-squared Euclidean distances as functions of the genomic distances between pairs of beads. In Fig. 7, we show the two boxplots for the highly and poorly expressed regions. A different behaviour is apparent: the poorly expressed region occupies less space than the highly expressed one, although their genomic spans are nearly the same. We checked the statistical significance of these features: by Kolmogorov-Smirnov tests, we can reject the hypothesis that the values from the two regions, for each genomic distance, are drawn from the same distributions. Interestingly, for genomic distances larger than 1.1 Mbp, the distributions of the mean-squared Euclidean distances are always leptokurtic. Moreover, beyond a genomic distance of 2 Mbp, the kurtoses from the highly expressed region are always larger than the ones related to the poorly-expressed region. This means that, statistically, the geometrical behaviour of the two stretches is similar for limited genomic distances but, in its entirety, the poorly expressed stretch is more folded than the other in most of its feasible configurations. These features can also be seen by inspecting visually our solutions, including the example configurations shown in Fig. 6, from which we observe that, besides being folded less tightly than the other, the highly expressed region very often interferes less with the rest of the chain, and is located at the periphery of the structure.

## Conclusion

We propose a multiresolution 3d chromatin structure estimation method based on sampling the solution space generated by a new score function made of a balanced mix of data-fit and geometrical requirements. The recursive multiresolution setting enables us to exploit the presence of nearly isolated genomic domains typically recurring at different scales. The geometrical constraints introduced implicitly in the score function allow our prior knowledge to be exploited coherently and flexibly through a small set of tuneable parameters. The solution space is sampled by simulated annealing, and the chromatin model is evolved through quaternion operators. The successive levels of genomic resolution, as well as the annealing parameters, are determined automatically.

The solutions obtained through an extensive experimentation performed on real hi-c data proved to reproduce the main features of the original contact frequency matrix, also avoiding excessive bending of the chromatin fibre and interpenetrations between beads. Besides being consistent with the data and physically plausible, our solutions show features that also suggest biological plausibility. In particular, the structures of two regions identified as highly and poorly expressed, respectively, are shown to possess the expected topological properties. As this feature was not specifically introduced in the model,^6^ we consider this result significant for a biological validation of our method, although this cannot be considered conclusive, and further experiments are needed to validate our algorithm biologically.

In summary, this approach offers algorithmic advantages over other approaches, as well as consistency and controllability features that make it a good candidate to generate and analyse large populations of structures fitting the same data. As a method, it is suitable to estimate the structure of any set of adjacent genomic fragments as soon as enough contact data are available. We made available online two versions of our code (one features a convenient GUI), written in Python and including all the classes and functions needed to run autonomously. Their routine use is straightforward, as only a few parameters need to be specifically set. The interested colleagues are invited to test it against different data sets and to suggest possible improvements or corrections, as well as to correlate their solution features to further known biological properties.

## Footnotes

1 http://www1.isti.cnr.it/˜salerno/ChromStructfolder.zip

3 The genomic sizes of our second-level beads are as follows (in Mbp, 0.7 being the minimum allowed size): 1.1; 2.2; 0.7; 2.1; 0.7; 1.8; 1.; 0.7; 0.7; 1.1; 2.; 1.6; 1.3; 0.7; 0.7; 1.8; 1.8; 1.7; 1.3; 0.8; 1.3; 0.9; 1.2.

3 Note that this threshold distance corresponds approximately to the largest distance penalised by the constraint term for the case *c* = 4, *b* = 5 in Fig. 2.

4 http://bioinfo.amc.uva.nl/HTMseq/controller(last accessed: 2016, June 15^*th*^).

5 http://www.ensembl.org/Release 77 (last accessed: 2016, June 15^*th*^).

6 No geometrical requirement introduced is site-specific.

## Acknowledgements

This work has been partially supported by the Italian Ministry of Education, University and Research, and by the National Research Council of Italy, Flagship Project InterOmics, PB.P05 (http://www.interomics.eu).

## Author’s contributions

CC conceived the modified bead-chain model and, with ES, wrote and debugged the code, run the experiments and evaluated the results. ES conceived the score function and some of the evaluation procedures and wrote the paper. MZ contributed in setting the initial parameters, suggested the validation strategy and participated in the evaluation of the results. AT, scientific responsible, Flagship Project InterOmics WP1-ISTI, suggested the research field and contributed statistical and algorithmic expertise and evaluated the results. All the authors have read and approved the manuscript.

## Availability of data and materials

The codes and data used to validate our method can be downloaded from http://www1.isti.cnr.it/˜salerno/ChromStructfolder.zip

## List of abbreviations

3C: Chromosome conformation capture
4C: Circularized chromosome conformation capture
5C: Carbon copy chromosome conformation capture
HCPC: Hierarchichal clustering on principal components
HiC: New generation sequencing technique introduced in [4]
GUI: Graphic user interface
M-S: Mean-squared
TAD: Topological association domain

## References

1. Dekker, J., Rippe, K., Dekker, M., Kleckner, N.: Capturing chromosome conformation. Science 295 (2002) 1306–1311

2. Zhao, Z.: Circular chromosome conformation capture (4c) uncovers extensive networks of epigenetically regulated intra- and interchromosomal interactions. Nature Genetics 38 (2006) 1341–1347

3. Dostie, J., Dekker, J.: Mapping networks of physical interactions between genomic elements using 5c technology. Nat. Protoc. 2 (2007) 988–1002

4. Lieberman-Aiden, E., van Berkum, N.L., Williams, L., Imakaev, M., Ragoczy, T., Telling, A., Amit, I., Lajoie, B.R., Sabo, P.J., Dorschner, M.O., Sandstrom, R., Bernstein, B., Bender, M.A., Groudine, M., Gnirke, A., Stamatoyannopoulos, J., Mirny, L.A., Lander, E.S., Dekker, J.: Comprehensive mapping of long-range interactions reveals folding principles of the human genome. Science 326 (2009) 289–293

5. van Berkum, N.L., Lieberman-Aiden, E., Williams, L., Imakaev, M., Gnirke, A., Mirny, L.A., Dekker, J., Lander, E.S.: Hi-c: a method to study the three-dimensional architecture of genomes. J. Vis. Exp. 39 (2010) 1869–1875

6. Varoquaux, N., Ferhat, A., Stafford Noble, W., Vert, J.P.: A statistical approach for inferring the 3d structure of the genome. Bioinformatics 30 (2014) i26–i33

7. Nowotny, J., Ahmed, S., Xu, L., Oluwadare, O., Hensley, N., Trieu, T., Cao, R., Cheng, J.: Iterative reconstruction of three-dimensional models of human chromosomes from chromosomal contact data. BMC Bioinf. 16 (2015) 338

8. Wang, S., Xu, J., Zeng, J.: Inferential modeling of 3d chromatin structure. Nucl. Ac. Res. 43 (2015) e54

9. Segal, M.R., Bengtsson, H.L.: Reconstruction of 3d genome architecture via a two-stage algorithm. BMC Bioinf. 16 (2015) 373

10. Trussart, M., Serra, F., Baù, D., Junier, I., Serrano, L., Marti-Renom, M.A.: Assessing the limits of restraint-based 3d modeling of genomes and genomic domains. Nucl. Ac. Res. 43 (2015) 3465–3477

11. Lajoie, B.R., Dekker, J., Kaplan, N.: The hitchhiker’s guide ti hi-c analysis: Practical guidelines. Methods 72 (2015) 65–75

12. Junier, I., Spill, Y.G., Marti-Renom, M.A., Beato, M., le Dily, F.: On the de-multiplexing of chromosome capture conformation data. FEBS Lett. 589 (2015) 3005–3013

13. Imakaev, M., Fudenberg, G., Mirny, L.A.: Modeling chromosomes: Beyond pretty pictures. FEBS Lett. 589 (2015) 3031–3036

14. Caudai, C., Salerno, E., Zoppè, M., Tonazzini, A.: Inferring 3d chromatin structure using a multiscale approach based on quaternions. BMC Bioinformatics 16 (2015) 234

15. Duggal, G., Patro, R., Sefer, E., Wang, H., Filippova, D., Khuller, S., Kingsford, C.: Resolving spatial inconsistencies in chromosome conformation measurements. Algorithms for Molecular Biology 8 (2013) 8

16. Caudai, C., Salerno, E., Zoppè, M., Tonazzini, A.: A statistical approach to infer 3d chomatin structure. In Zazzu, V., et al., eds.: Mathematical Models in Biology. Springer International Publishing Switzerland, Cham (2015) 161–171

17. Dixon, J.R., Selvaraj, S., Yue, F., Kim, A., Li, Y., Shen, Y., Hu, M., Liu, J.S., Ren, B.: Topological domains in mammalian genomes identified by analysis of chromatin interactions. Nature 485 (2012) 376–380

18. Vince, J.A.: Geometric Algebra for Computer Graphics. Springer, Berlin (2008)

19. Kirkpatrick, S., Gellatt, C.D.J., Vecchi, M.P.: Optimization by simulated annealing. Science 229 (1983) 671–680

20. Thode, H.C., Jr.: Testing for Normality. Marcel Dekker, New York (2002)

21. Mateos-Langerak, J., Bohn, M., de Leeuw, W., Giromus, O., Manders, E.M.M., Verschure, P.J., Indemans, M.H.G., Gierman, H.J., Heermann, D.W., van Driel, R., Goetze, S.: Spatially confined folding of chromatin in the interphase nucleus. PNAS 106 (2009) 3812–3817

22. Husson, F., Josse, J., Pagès, J.: Principal component methods - hierarchical clustering - partitional clustering: why would we need to choose for visualising data? Technical report, Applied Mathematics Department, Agrocampus, Rennes, France (September 2010)

23. Versteeg, R., van Schaik, B.D.C., van Batenburg, M.F., Roos, M., Monajemi, R., Caron, H., Bussemaker, H.J., van Kampen, A.H.C.: The human transcriptome map reveals extremes in gene density, intron length, gc content, and repeat pattern for domains of highly and weakly expressed genes. Genome Res. 13 (2003) 1998–2004

